# The role of FRUITFULL controlling cell cycle during early flower development revealed by time series snRNA-seq experiments

**DOI:** 10.1101/2025.08.31.673337

**Authors:** Peilin Chen, Xiaocai Xu, Cezary Smaczniak, Bénédicte Desvoyes, Crisanto Gutiérrez, Robert Sablowski, Kerstin Kaufmann, Jose M Muino

**Affiliations:** Plant Cell and Molecular Biology, Institute of Biology, Humboldt-Universität zu Berlin, Berlin, Germany; Centro de Biología Molecular Severo Ochoa, CSIC-UAM, Cantoblanco, Madrid, Spain; Cell and Developmental Biology Department, John Innes Centre, Norwich Research Park, Norwich NR4 7UH, United Kingdom; German Federal Institute for Risk Assessment (BfR), German Centre for the Protection of Laboratory Animals (Bf3R), Max-Dohrn-Straße 8-10, 10589 Berlin, Germany

**Author notes:** Contributed equally (Peilin Chen, Xiaocai Xu).

**Keywords:** Flower development, cell cycle, scRNA-seq analysis, pseudo-time inference

## Abstract

**BACKGROUND:** Starting from pools of undifferentiated cells, plants generate new organs post-embryonically in response to external and endogenous signals. This requires a dynamic coordination of cell division with cellular growth and differentiation regulatory programs. However, little is known how this coordination is achieved at the molecular level during flower development.

**RESULTS:** We used time-series single-nucleus RNA sequencing (snRNA-seq) experiments of synchronized *Arabidopsis thaliana* flower developmental stages to characterize the transcriptome dynamics and the connections between cell cycle and developmental regulatory programs during early flower development. The results show a bifurcation between transcriptional trajectories corresponding to cell cycle progression and floral development. We identify the regulation of the cell cycle inhibitor KIP-RELATED PROTEIN 2 (KRP2) by FRUITFULL (FUL) as a key regulatory point on this bifurcation point, and validate the importance of this regulation *in vivo*.

**CONCLUSIONS:** Our work illustrates how time-series snRNA-seq experiments can be used to identify bifurcation points between regulatory programs and to identify candidate regulators on these bifurcations. In particular, we identify the regulation of KRP2 by FUL as an important regulatory point to balance cell division and developmental differentiation in plants.

## Background

Organismic growth and development are complex and highly regulated processes. Cell proliferation and cell differentiation needs to be balanced in a stage- and cell-type specific fashion ^1^. This is even more important in higher plants like *Arabidopsis thaliana,* where organs are formed responding to endogenous and environmental stimuli during postembryonic development. Therefore, plants need regulatory systems to replenish and maintain their pool of pluripotency cells in the meristems ^2^, and, responding to local stimuli, to initiate cellular growth and differentiation programmes to form mature organs and tissues. Terminal differentiation is often associated with cell cycle exit, but cell fate decisions are frequently linked to cell cycle transitions ^3^. In human pluripotent stem cells the transition through mitosis and G1 is essential to exit pluripotency and enter differentiation ^3^. However, mouse hematopoietic stem cells are primed to differentiate to megakaryocytes in the G2 phase by a mechanism involving DNA damage ^4^. This is, cell fate is determined at different stages of the cell cycle in multipotent cells, however it is not clearly understood how this happens in plant multipotent cells.

Cyclin-dependent kinases (CDKs) are main regulators controlling transition through the different cell cycle phases by phosphorylation of many substrates ^5^. A-type CDKs are essential at both G1-to-S and G2-to-M transitions, while B-type CDKs display maximum activity during the G2-to-M transition and the M-phase ^6^. CDK activity is largely influenced by their association with different cyclin patterns. For example, D-type cyclins mainly regulate G1-to-S transition, whereas A-type and B-type cyclins control S-phase progression and G2-to-M transition respectively ^6^. These processes include the binding of small CDK inhibitory proteins, such as KIP-RELATED PROTEINs (KRPs), and the control of cyclin proteolysis by the anaphase-promoter complexes ^6^. During Arabidopsis development, KRPs show distinct expression profiles both spatially and temporally ^7^. For example, in the shoot apical meristem (SAM), KRP1 and KRP2 express weakly in the central meristem, and the expression of KRP2 is comparably abundant in deep rib meristem. On the contrary, the expression of KRP3 and KRP5 are mainly in the peripheral zone. As it reaches the highest abundance in floral buds ^7,8^, *KRP2* is down-regulated by DELLA proteins to maintain the inflorescence meristem (IM) size ^9^, and repressed by JAGGED (JAG) transcription factor to constrain floral organ morphogenesis ^10^. High expression levels of KRP2 in transgenic plants hinder both DNA replication and mitosis ^7,11^ , indicating that KRP2 overexpression inhibits both G1-S and G2-M transition. KRP2 overexpression driven by the meristematic-specific *SHOOT MERISTEMLESS* gene leads to an increase in the ploidy of the cells, indicating that KRP2 overexpression can be associated with the onset of endoreduplication. In Arabidopsis, endoreduplication in leaf cells is regulated among others by KRP2 ^12^. In addition, KRP2 is reported to inhibit the G1 to S transition in lateral root initiation ^13^. However, we still lack a more comprehensive understanding on regulatory mechanisms controlling cell cycle plant transitions depending on developmental cues ^3,4,14^.

One of the main regulators of plant development is the MADS-box transcription factor (TF) family 15. In particular, during the early stages of flower development, the AP1/FUL subfamily plays important roles. For example, APETALA1 (AP1) promotes the transition from IM to floral meristem (FM) (Kaufmann et al., 2010b). FRUITFULL (FUL), a paralog of AP1, plays important regulatory roles during the vegetative to reproductive transition of the shoot apical meristem by promoting flowering time (Ferrándiz et al., 2000a). It does this by repressing *FLOWERING LOCUS C* (*FLC*) and competing with FLC for common targets like *SUPPRESSOR OF OVEREXPRESSION OF CO 1 (SOC1)*. During the transition from IM to FM, FUL directly binds and enhances SOC1 expression, which then partners with FUL to activate *LEAFY* (*LFY*) and repress *TERMINAL*

*FLOWER1* (*TFL1*) 16,17. Additionally, FUL controls IM fate by repressing *APETALA2* (*AP2*) and related genes, thereby influencing SAM activity 18,19. Interestingly, recent reports indicate that FUL can affect the cell cycle, in particular, *ful-1* mutants show higher division activity in SAMs 20. However, it is not clear how FUL may influence the cell cycle in the SAM.

Single cell genomics technologies enable us to study the dynamic trajectories of gene activities during cellular growth and differentiation. In mammals, these technologies have been successfully used to study cell cycle progression and regulation (e.g. ^21,22^). Therefore, we decided to study the gene expression dynamics during the early stages of flower development of *Arabidopsis thaliana*, with a special focus on cell cycle progression and regulation in meristematic cells.

## Results

### snRNA-seq time series of the early stages of flower development

To obtain a high-resolution atlas of gene expression dynamics in early Arabidopsis flower development, we performed single nuclei RNA-seq (snRNA-seq) experiments using a system for synchronized floral induction (*pAP1:AP1-GR ap1-1 cal-1)* ^23^. This system allows us to collect plant material from specific flower developmental stages to be characterized with snRNA-seq experiments. In particular, we collected inflorescence/flower tissues after 0 (uninduced), 2, 4, and 8 days of Dexamethasone (DEX) induction which roughly corresponds to stages 0-9 of flower development as defined by Ryan *et al*. (2015) and Smyth *et al.* (1990)^23,24^, or stages F19+ to F12 as defined by TraVA ^25,26^ (Figure 1a). Thus, we cover early stages of meristem patterning to organ differentiation. In this study, we will refer to these samples as S0, S2, S4, and S8. We chose these particular stages because they represent key time points in early flower development, and we previously characterized them using other genomics approaches ^27,28^. Additionally, we reanalyzed snRNA-seq ^29^ of wild-type *Arabidopsis* flower tissues (mix tissues before anthesis, until stage 12 ^24^), to be used as a scaffold to integrate the others snRNA-seq datasets. We call this sample Col0.

**Figure 1.**
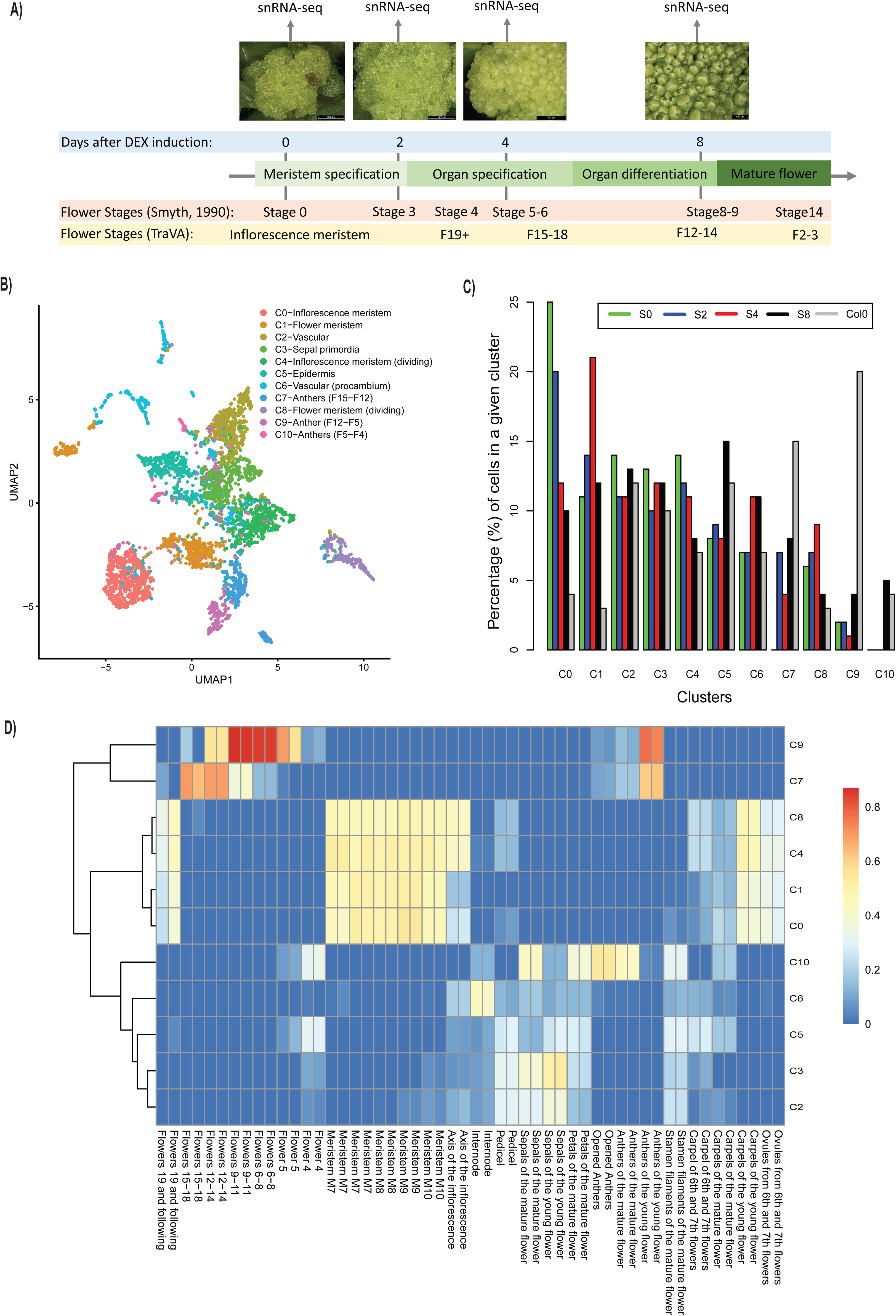
Analysis and annotation of the time series snRNA-seq experiments. **A)** Schematic representation of the developmental time points used to generate snRNA-seq datasets. At the bottom, it is shown the correspondence between the time points used and the developmental flower stages defined by Smyth *et al.*, 1990 ^24^, and Klepikova *et al.*, 2015 and 2016^25,26^. **B)** UMAP plot and cluster annotation of the combined snRNA-seq analysis. **C)** Bar plot showing the proportion of transcriptomes from each snRNA-seq developmental stage assigned to each identified cluster. **D)** Heatmap showing the Pearson correlation between the average expression of each cluster (rows) and selected bulk RNA-seq datasets from TraVA (columns). To calculate the correlation only genes identified as top 50 marker genes for each cluster were used.

SnRNA-seq libraries were prepared using the nanowell-based ICELL-8 system from Takara as previously described ^29^. We obtained ∼4,500 transcriptomes among all samples, with an average of ∼4,200 genes detected and 2*10^5^ mapped reads per transcriptome (Supplementary Figure S1a-b). These numbers are typical for the ICELL-8 system. The low number of reads mapping to mitochondrial genes (<5%) indicates low organelle contamination (Supplementary Figure S1d). Our snRNA-seq can recapitulate the gene expression patterns of bulk RNA-seq ^28^ data from the same stage and tissue samples (Supplementary Figure S2; median Pearson correlation of R=0.76). Integrative analysis with Liger v0.4.2 identified 11 clusters across all samples (Figure 1b). To annotate the clusters, we identified the top 50 marker genes for each cluster (Supplementary Table S1) and correlated their average expression in each cluster against publicly available bulk RNA-seq datasets of different plant tissues (Figure 1d).

Clusters C0, C1, C4, and C8 represent meristematic tissues as indicated by their strongest correlation with bulk RNA-seq profiles of meristematic tissues from TraVA (Figure 1d). Clusters C0 and C4 seem to be at the early stage of inflorescence meristems as these clusters are enriched in the uninduced snRNA-seq sample (S0), representing ∼25% and ∼15% of transcriptomes of this sample (Figure 1C). Examples of markers of cluster C0 (Supplementary Table S1) are *AT-HOOK MOTIF NUCLEAR-LOCALIZED PROTEIN 15* (*AHL15*), a regulator of *FUL* and *SOC1* ^30^, and *MULTICOPY SUPPRESSOR OF IRA1 4* (*FVE*) one of the autonomous pathways controlling vegetative-flower transition by negatively regulating *FLOWERING LOCUS C (FLC)* expression ^31^ (Supplementary Figure S3a). C4 is likely meristematic dividing cells given the large proportion of marker genes involved in cell division. For example, *PHRAGMOPLAST ORIENTING KINESIN 1* and *2* (*POK1* and *POK2*) encoding kinases involved in cytokinesis ^32^, and *ARABIDOPSIS THALIANA KINESIN 1* and *5* (*ATK1* and *5*) encoding kinases involved in spindle morphogenesis ^33^ (Supplementary Figure S3a). Cluster C1 and C8 are likely meristematic cells from later stages as they are enriched in the samples collected after 4 days of induction (∼20%, and 10% of the transcriptomes of this sample, respectively. See Figure 1c). This stage corresponds with the time of flower organ specification; therefore, these two clusters are likely enriched in flower meristems. Examples of marker genes from cluster C1 are: *ABNORMAL FLORAL ORGANS* (*AFO*), encoding a YABBY transcription factor expressed in floral organ primordia ^34^, and *NIMA-RELATED KINASE 2* (*NEK2*), a kinase gene expressed in young leaves and apical buds involved in the cell-cycle regulation ^35^. Marker genes of cluster C8 were strongly enriched in genes involved in cell division. For example, *CELL DIVISION CYCLE 20.1 and 20.2* and *MITOTIC ARREST-DEFICIENT 2* (*MAD2*) encode proteins that are part of the mitotic checkpoint complex ^36^ (Supplementary Figure S3a).

Clusters C7, C9, and C10 seem to represent transcriptomes from anthers as they show the strongest correlation with bulk RNA-seq from these tissues (Figure 1b). Cluster C7 is enriched in earlier flower stages than C9 as shown by the correlation with the time-series RNA-seq data from TraVA (Figure 1d), and C10 seems to represent anthers from the mature flower (Figure 1d). Examples of marker genes of C7 are *TAPETUM-SPECIFIC METHYLTRANSFERASE 1* (*TSM1*), encoding a methyltransferase expressed exclusively in the tapetum of developing stamens ^37^, and *MALE STERILITY 1* (*MS1*), encoding a PHD-type transcription factor regulating pollen and tapetum development ^38^. Examples of C9 marker genes are *ABORTED MICROSPORES* (*AMS*), a basic helix-loop-helix (bHLH) transcription factor gene regulating tapetal cell development and pollen cell wall ^39^, and *CYTOCHROME P450* (*CYP704B1*), a cytochrome gene expressed during the development of anthers and involved in the creation of the exine in the pollen wall ^40^. Examples of markers from cluster C10 are: *DIHYDROFLAVONOL 4-REDUCTASE-LIKE1* (*DRL1*), encoding an oxidoreductase required for pollen development and male fertility ^41^, and *LESS ADHERENT POLLEN 3* (*LAP3*), a gene involved in pollen development and exine structuring ^42^ (Supplementary Figure S3a).

Cluster C6 seems to be enriched in vascular tissues (Supplementary Figure S3b), in particular procambium. Examples of marker genes are *PHLOEM INTERCALATED WITH XYLEM* (*PXY*), a receptor-like kinase gene expressed in the procambium which represses its differentiation ^43^, and *HOMEOBOX GENE 8* (*ATHB-8*), encoding a bHLH transcription factor regulating procambial cell fate acquisition ^44^ (Supplementary Figure S3a). Cluster 2 also seems to be vascular tissues because of its strong correlation with the S17 bulk RNA-seq data (Supplementary Figure S3b). S17 is a marker of phloem cells ^45^. Few marker genes were reliably detected in this cluster (Supplementary Table S1).

Cluster C5 seems to be enriched in epidermal cells (Supplementary Figure S3b). Examples of marker genes in this cluster are: *PROTODERMAL FACTOR 2* (*PDF2*) and *1* (*PDF1*), encoding homeodomain transcription factors that play a major role in maintaining L1 epidermic cell identity, and *LONG-CHAIN ACYL-COA SYNTHASE 1* (*LACS1*) and *2* (*LACS2*), acyl-CoA synthetases involved in cuticular wax and cutin biosynthesis ^46^ (Supplementary Figure S3a).

Finally, cluster C3 seems to be enriched in sepal tissues, in particular photosynthetic mesophyll cells, because of the strong correlation with bulk RNA-seq of sepals of the young flower from TraVA (Figure 1d). Examples of markers for these clusters are related to photosynthesis activity: *PHOTOSYNTHETIC NDH SUBCOMPLEX L 4* (*PNSL4*), a gene encoding a member of the NDH complex ^47^, and *CHLORORESPIRATORY REDUCTION 3* (*CRR3*) which is essential to stabilize the NDH complex ^48^ (Supplementary Figure S3a).

In summary, we have generated a time-series snRNA-seq experiment of the early stages of flower development and annotated the main clusters identified on this dataset. In particular, we have identified four clusters representing meristematic tissues to be studied in the next sections. The synchronized system used allowed us to locate the obtained transcriptomes in a developmental coordinate system and to enrich early meristematic stages that other ways will be too transient to be captured (Figure 1c).

### Reclustering of meristematic cells identifies a bifurcation point between cell cycle and developmental programs

Since we were able to identify the transcriptome clusters corresponding to meristematic tissues (C0, C1, C4, and C8), we next aimed to characterize the transcriptomic developmental dynamics of inflorescence and floral meristems. For this, we performed a pseudo-time analysis of the subset of transcriptomes belonging to these clusters using Monocle3 ^49^. To have an outgroup to facilitate the new clustering, the transcriptomes from cluster C3 were also included in the pseudo-time analysis as they represent transcriptomes from young sepal mesophyll tissues (Figure 1d). Sepals are the first floral organs to grow from floral meristems, and this process starts as early as stage 2 of flower development ^24^. The resulting UMAP plot and reclustering is shown in Figure 2a. Two clear trajectories are observed, with a bifurcation point starting at cluster 2. Cluster 6 seems to represent the earliest developmental stage as it is enriched in the uninduced snRNA-seq sample (S0, see Figure 2b). The marker genes of the clusters from one of the two trajectories are enriched in genes related to the cell cycle (Supplementary Table S2), for example: *CELL DIVISION CYCLE 20.1 and 2* (*CDC20.1-2*), *FIZZY-RELATED 3* (*FZR3*), *INFLORESCENCE MERISTEM RECEPTOR-LIKE KINASE 2* (*IMK2*), and *CYCLIN B2;4* (*CYCB2;4*), the other trajectory seems to be related with a flower-specific developmental program as it is enriched in flower developmental genes, for example: *SEPALLATA3* (*SEP3*), *ULTRAPETALA1* (*ULT1*) or *AINTEGUMENTA-LIKE 5* (*AIL5*). Figure 2c shows the expression of several known cell cycle marker genes across the detected clusters of the cell cycle trajectory. We used this information to approximately locate the different stages of the cell cycle in Figure 2a. Interestingly, the order of the clusters in the UMAP plot follows the expected order of the cell cycle, this is G1->S->G2->M. The bifurcation point of the pathways is mapped to cluster 2 which should correspond to the S phase. We notice that, at this moment, we cannot separate the mitotic from the endoreplication cell cycle as some cell cycle stages are common to both. The second pathway found in the UMAP (Figure 2A) can be related to developmental time as shown by their expression correlation with bulk-RNA-seq time-series experiments from TraVA (Figure 2D) and the distribution of snRNA-seq samples collected at different developmental times across the UMAP plot (Figure 2C, Supplementary Figure S4). A closer inspection of the co-expression of the clusters in this pathway to the different TraVa flower expression profiles reveals that the first half of the clusters on the developmental-related trajectory ordered by their pseudo-time (Clusters 6, 2, 10, 15, 13, and 4), likely, represent meristematic tissues and the other clusters (Clusters: 12, 1 18, 9 and 14) mainly represent sepal-related tissues (Supplementary Figure S4G), as expected by the fact that we have pre-selected transcriptomes enriched in meristems and sepals tissues from the initial dataset (Figure 1B). Clusters 12 and 18 are annotated as anther tissues which indicates that the annotation of these two clusters were not accurate, or that our pre-selection of meristematic cells contains a low percentage of non-desired transcriptomes. However, they are located far from the branching point, and it is unlikely to have any effect on any further study of the expression around the branching point.

**Figure 2.**
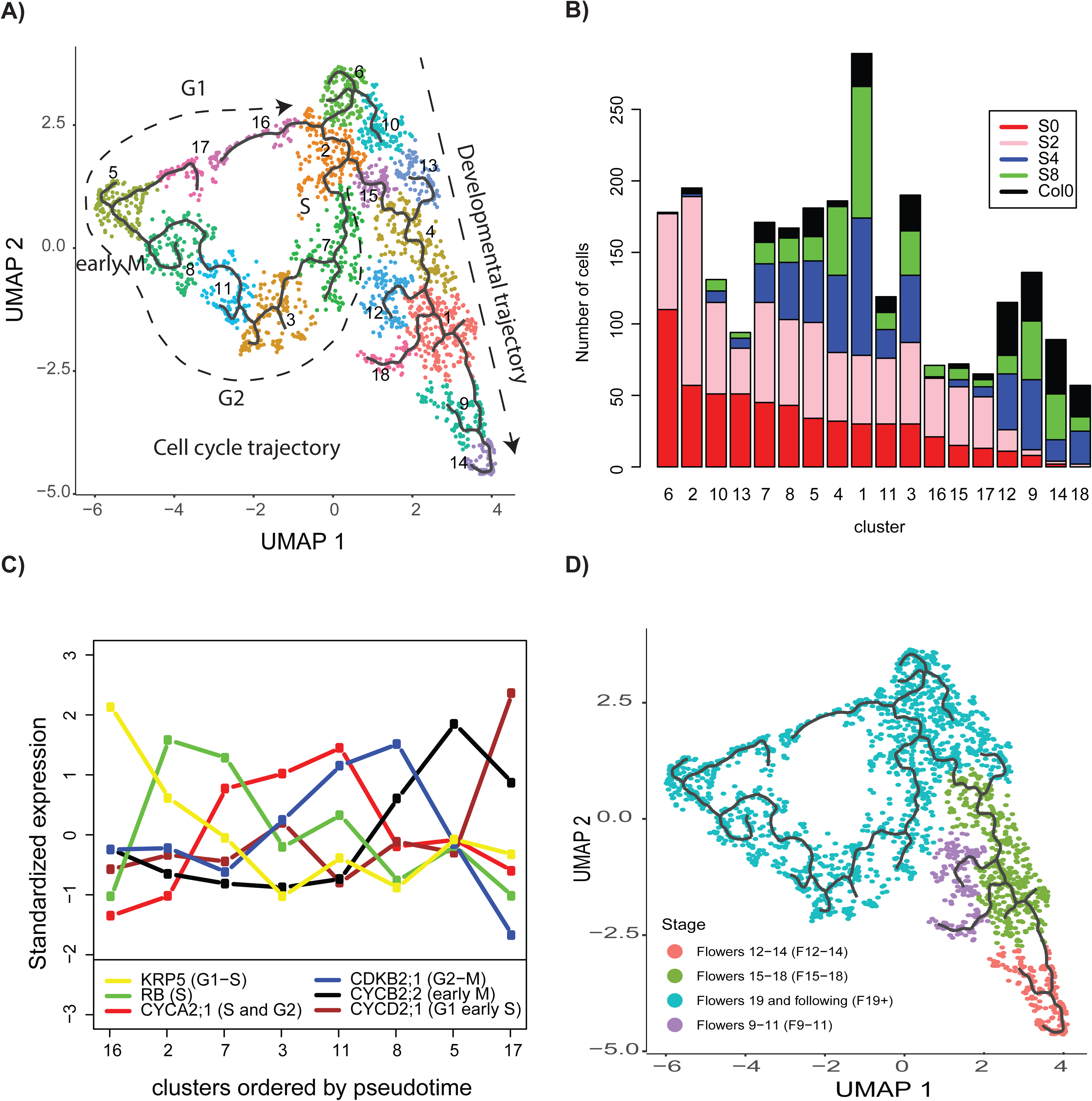
Analysis of the bifurcation point between cell cycle and cell developmental programs. **A)** Reclustering of cells belonging to meristematic and sepal primordia clusters (C0, C1, C3, C4, and C8) identified in Figure 1a were reanalyzed and clustered using Monocle3. Continuous black line indicates the estimated pseudo-time trajectory. Two main branches were detected and indicated by the dashed arrows. For the cell cycle branch, the approximate location of the different cell cycle phases is indicated based on Figure 2C. **B)** The bar plot shows the number of transcriptomes (y-axis) per snRNA-seq sample in each cluster defined in A (x-axis). The x-axis is decreasingly ordered by the number of transcriptomes of the uninduced sample (S0). This, roughly, orders the clusters from earlier (left side) to later stages (right side). **C)** Expression of known cell cycle marker genes in selected clusters (x-axis) identified in A. **D)** Annotation of each cluster depending on its expression correlation with bulk RNA-seq (TraVa, time-series). The correlation was calculated between the average gene expression for each cluster and TraVA developmental stage-specific bulk RNA-seq data. The TraVa dataset with the highest correlation is reported for each cluster.

Next, we estimate the pseudo-time value for each transcriptome with Monocle3. We used cluster 6 as the initial point as it is enriched in the uninduced sample (S0, Figure 2b). For the branch belonging to the developmental-related trajectory, we observed that the pseudo-time values assigned to the transcriptomes of each snRNA-seq sample (S0, S2, S4, and S8, see Supplementary Figure S5A, B) are linked with the developmental stage when the sample was collected. SnRNA-seq samples from early developmental time points have lower pseudotime values than samples collected at later developmental time points, reinforcing the idea that this branch is related to a developmental trajectory. On the other hand, for the branch related to the cell cycle, the samples collected at different time points after DEX induction have no difference in pseudo-time values distribution, suggesting that the estimated pseudo-time on this branch is not dependent on the developmental stage of the meristem. Indeed, we show in Figure 2C that the different clusters of the cell cycle branch are related to the different cell cycle phases, therefore, the pseudo-time estimation in this cell population is related to cell cycle time rather than developmental time. Next, we aimed to identify potential genes that are associated with this bifurcation as a first step to identifying regulators that control this very important step.

Monocle3 offers the possibility to identify genes whose expression is associated with a bifurcation point through the Moran’s I test. To find cell cycle genes associated with this bifurcation point, we performed the Morańs I test using the expression of core cell cycle genes defined by ^50^. Only *KIP-RELATED PROTEIN 2* (*KRP2*) was significant at an FDR level <0.05. KRP2 is a cyclin-dependent kinase inhibitor that negatively regulates cell cycle transitions (G1/S, and G2/M) ^12^. Other genes may be linked to this bifurcation point, but we may not detect them given the properties of our snRNA-seq experiment. Notably, low-expressed genes will not likely be consistently detected in the snRNA-seq experiment, so other important cell cycle factors could be missed from this analysis.

The average expression of the *KRP2* gene in individual clusters shows that it has the highest expression in cluster 15, in the development trajectory just after the bifurcation point (Supplementary Figure 4F-G). It has relatively low expression in cluster 2, where the bifurcation point is located, and cluster 7, in the cell-cycle trajectory just after the bifurcation point (Supplementary Figure 4G-H). This indicates that an increased expression of *KRP2* may be associated with the controlling cell cycle transition in cluster 15. Next, we wanted to identify the transcription factor (TF) regulators controlling *KRP2* expression in cluster 15. Therefore, we used GENIE3 ^51^ to estimate the gene regulatory network of *KRP2* on this cluster (Supplementary Table S3). The estimated top 5 transcription factor regulators of *KRP2* in cluster 15 were AT5G18550, SEPALLATA2 (SEP2), ETHYLENE RESPONSE FACTOR 74 (ERF74), KANADI 2 (KAN2), and FRUITFULL (FUL). AT5G18550 is a Zinc finger protein with little information publicly available. SEP2 is a MADS-box TF associated with flower meristem determinacy ^52^ and development ^15^, and it has been reported to form protein complexes with FUL *in vivo* ^17^. ERF74 is an ethylene response factor of the AP2 transcription factor family involved in sensing of reactive oxygen species (ROS) ^53^. Recent reports identify ROS signaling as an important factor in meristem maintenance and mitosis ^54^. KAN2 is a member of the KANADI TF family and has been reported to affect cell division patterns ^55^. Finally, FUL is a MADS-box TF associated with inflorescence and flower development ^15^, and it has been linked to cell cycle regulation since the number of dividing cells increased in *ful* mutant plants ^20^.

In a previous publication, we characterized the FUL DNA binding landscape in inflorescence meristems by ChIP-seq experiments ^17^. We identified that FUL can bind to a region ∼1.3 kb downstream of the *KRP2* locus (Figure 3A, and Supplementary Table 3b of van Mourik *et al.* (2023)^17^. This region has been described as a *KRP2* regulatory region bound by JAGGED and REPRESSOR OF GA (RGA1) TFs ^9,10^. Bulk RNA-seq experiments in inflorescence meristems ^17^ showed that *KRP2* decreases in expression in the *ful-7* mutant compared to WT (*P*< 0.003). This indicates that FUL is a potential activator of *KRP2* in this tissue. Interestingly, FUL seems to not only regulate *KRP2*, but also regulate other genes encoding KRPs and KRP regulators. In our previous ChIP-seq experiments, we found that FUL has binding sites in the 3 kb upstream to 1 kb downstream regions of all *KRP* genes except *KRP5* (see Supplementary Figure S6, Supplementary Table S4). This reinforces the idea that FUL may regulate this pathway. FUL also binds to the DELLA genes *RGA1* and *RGA-LIKE 2* (*RGL2*) involved in the regulation of *KRP*s ^9^. *KRP7* (*P*< 0.04) and the DELLA gene *RGL2* (*P*<0.001) increase in expression in the *ful-7* mutant compared to WT (Supplementary Table S4). Therefore, FUL is a potential repressor of these genes in this tissue. *KRP1* and *KRP6* had too low expression in our bulk RNA-seq data to be able to test any change in expression. The other *KRPs* don’t show any significant change in expression. However, *KRPs* are usually expressed in specific cells, and our bulk RNA-seq experiments may not be able to identify change in expression in bulk tissues.

**Figure 3.**
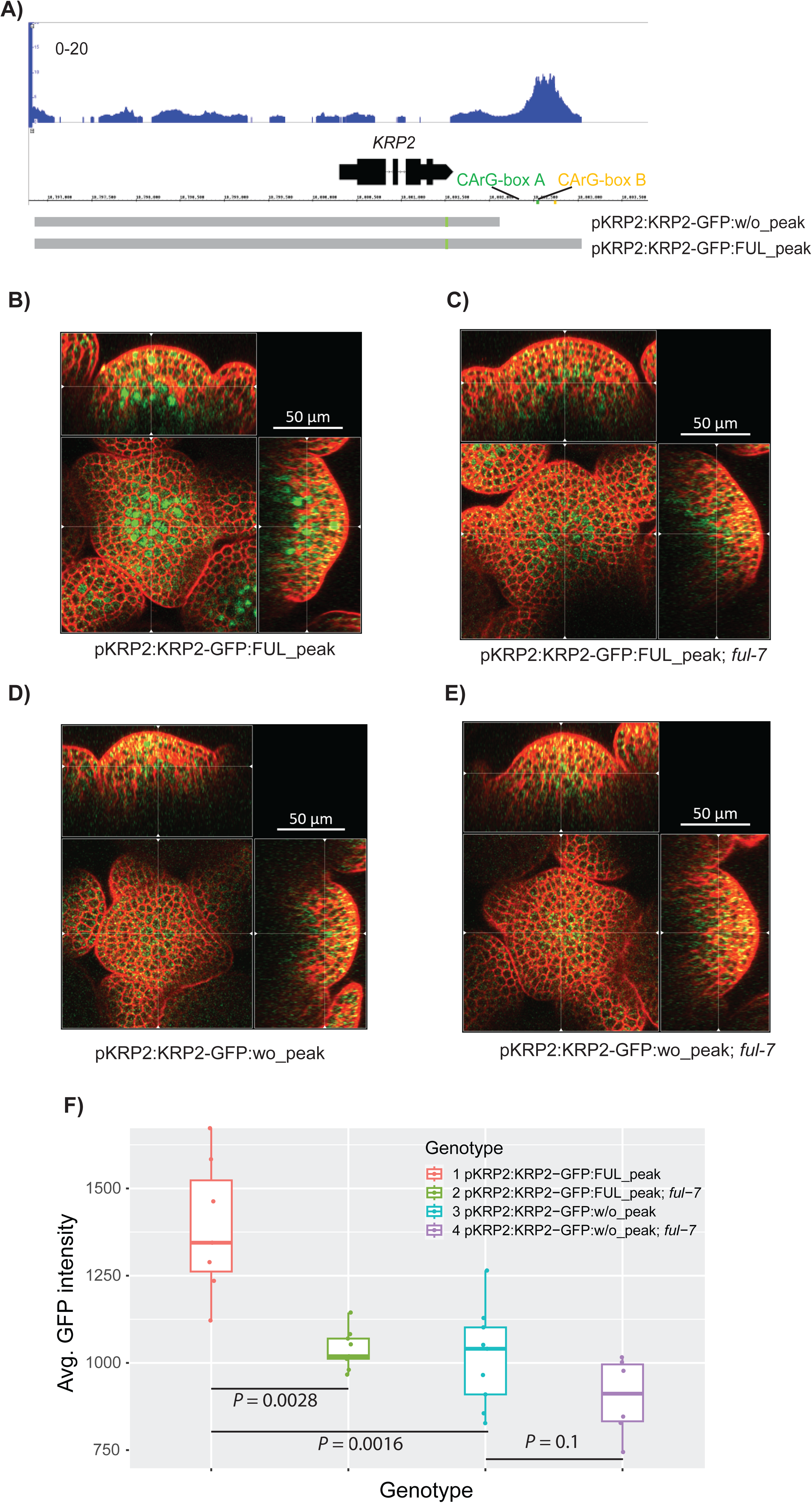
Confocal analysis of KRP2-GFP localization in inflorescence meristem. **A)** Screenshot of FUL ChIP-seq binding peak near *KRP2* gene. CArG-box A and B indicate the position of the only two CArG-box motifs located under the FUL ChIP-seq peak. The genomic region of *KRP2* in the reported construct KRP2p:KRP2-GFP (Serrano-Mislata et al., 2017) was shown below the screenshot. pKRP2:KRP2-GFP:w/o_peak and pKRP2:KRP2-GFP:FUL_peak indicate the genomic regions used for GFP reporter constructs cloning, one without the FUL ChIP peak region and the other contains the FUL peak region. **B-E)** Confocal images of IMs with ortho views. The genotypes are: (B) *pKRP2:KRP2-GFP:FUL_peak* (N = 7), (C) *pKRP2:KRP2-GFP:FUL_peak* in *ful-7* (N = 9), (D) *pKRP2:KRP2-GFP:w/o_peak* (N = 9), E) *pKRP2:KRP2-GFP:w/o_peak* in *ful-7* (N = 6). The plants used for scanning are at a similar age with only 1-3 open flowers. **F)** Average GFP intensity of inflorescence meristem cells. *P* value was calculated using bilateral t-test.

In summary, we have identified a bifurcation point between cell cycle and developmental-related programs during early flower development by using time-series snRNA-seq experiments. Our bioinformatic analysis estimates that *KRP2* activity is linked to this bifurcation point, and predicts several important regulators affecting the *KRP2* expression in this process. The regulation of *KRP2* by FUL seems a particularly interesting hypothesis as we have previous experimental evidence supporting this regulation.

### FUL increases KRP2 expression by binding in the *KRP2* downstream region

Next, we aimed to validate the regulation of *KRP2* by FUL *in planta*. The potential regulatory region downstream of *KRP2* (Figure 3A) bound by FUL contains two CArG-box motifs, which are the sequences expected to be bound by FUL. Our EMSA experiments showed that FUL can bind *in vitro* to each of these two individual CArG-boxes as a homodimer or heterodimer (Supplementary Figure S7). This observation narrows down the location of the potential genomic regulatory region controlled by FUL. Previous study ^9^ has characterized *KRP2* expression using a construct that covers only one of these two CArG-boxes. In this study, we generated two new constructs to study *KRP2* expression (Figure 3A), one containing both CArG-boxes (pKRP2:KRP2-GFP:FUL_peak) and another excluding both CArG-boxes (pKRP2:KRP2-GFP:w/o_peak). These constructs were used to transform *Col*-0 WT, and *Col*-0 *ful-7* plants.

FUL has been reported to be expressed in the nucleus of young flowers, especially in the inflorescence and flower meristems ^56^. In the inflorescence meristem, FUL:GFP signal has been observed in all cell layers. During the initial stages of the flower bud primordium development the signal starts to decrease in the subepidermal and inner cell layers. ^9^ has reported that their pKRP2:KRP2-GFP construct is expressed weakly below the inflorescence and floral meristems. Our pKRP2:KRP2-GFP:FUL_peak containing the two CArG-boxes recognized by FUL has a stronger expression, but it is also mainly restricted to the inner cell layers of the inflorescence meristem (Figure 3B) and flower meristem (Supplementary Figure S8-9). This overlap in expression is in line with the hypothesis that FUL can regulate *KRP2*. In the *ful-7* mutant background, the signal of pKRP2:KRP2-GFP:FUL_peak significantly decreases (bilateral t-test, P< 0.0029 Figure 3C, F), indicating that FUL is essential for *KRP2* expression in inflorescence meristems. There is also a similar decrease in *KRP2* expression compared to WT (bilateral t-test, P< 0.0016 Figure 3D, F) when using pKRP2:KRP2-GFP:w/o_peak in the WT background. This is, the elimination of the genomic region containing the two CArG-box motifs bound by FUL has a similar effect on the expression of *KRP2* as eliminating *FUL* expression. A similar pattern is observed in flower meristems (Supplementary. Figure S8-S9). This supports the hypothesis that FUL activates *KRP2* by binding to these two CArG-box regions.

In summary, we showed that FUL regulates the expression of the cell cycle regulator *KRP2* in IM and flower meristems, and that this depends on the presence of the genomic regulatory region downstream of *KRP2* that is bound by FUL. This strongly supports the hypothesis that FUL directly upregulates *KRP2* upon binding on its downstream regulatory region.

### FUL participates in the cell cycle progression

As mentioned above, *KRP2* is associated with cell cycle transition inhibition, and FUL plays a positive role in the transcriptional regulation of *KRP2.* This regulation is predicted to happen after cluster 2 (Figure 2a) which corresponds to the S-phase, where KRP2 protein will start to accumulate. This KRP2 accumulation may play an important role controlling the G2 to M transition. To experimentally validate the effect of FUL in the G2 to M transition, we introduced the Plant Cell Cycle Indicator (PlaCCI) construct ^57^ into Col-0 WT, and Col-0 *ful-7* mutant plants (Figure 4 A, B), and checked the cell cycle proportion changes between these two lines using young IMs when there are one to three flowers open (within one week after bolting). The PlaCCI construct has a single cassette containing pHTR13::HTR13-mCherry, pCDT1a::CDT1a-eCFP, and pCYCB1;1::NCYCB1;1-YFP reporters that can indicate different cell cycle phases respectively. After splitting the channels in ImageJ and analyzing them by MorphoGraphX 2.0 software (MGX), we obtained the numbers of nuclei expressing the corresponding reporter for each channel and calculated the proportion of the channel-specific nuclei versus total nuclei (Figure 4 C). Compared to WT, *ful-7* mutant plants have a significantly lower proportion of cells expressing H3.1-mCherry which is the S and early G2 phase reporter gene of the PlaCCl system (two-tailed *t*-test, *P*=0.0169), and a higher proportion of cells accumulating G1 phase reporter CDT1a-CFP (two-tailed *t*-test, *P*=0.0469). We didn’t detect a significant change in the proportion of cells expressing CYCB1;1-YFP, the reporter of late G2 and M phases, this is likely due to the low number of nuclei expressing CYCB1;1-YFP, as expected by the short length of the mitotic (M) phase (Desvoyes et al., 2020). The decrease of G2 cells in *ful-7* isline with the hypothesis that the activation of KRP2 by FUL is required to block G2 to M transition in IM. When FUL is present, cells can be blocked at G2 phase and therefore there will be a higher accumulation of G2 cells than when in *ful-7*. This shows that FUL plays an important role controlling cell cycle progression in the IM.

**Figure 4.**
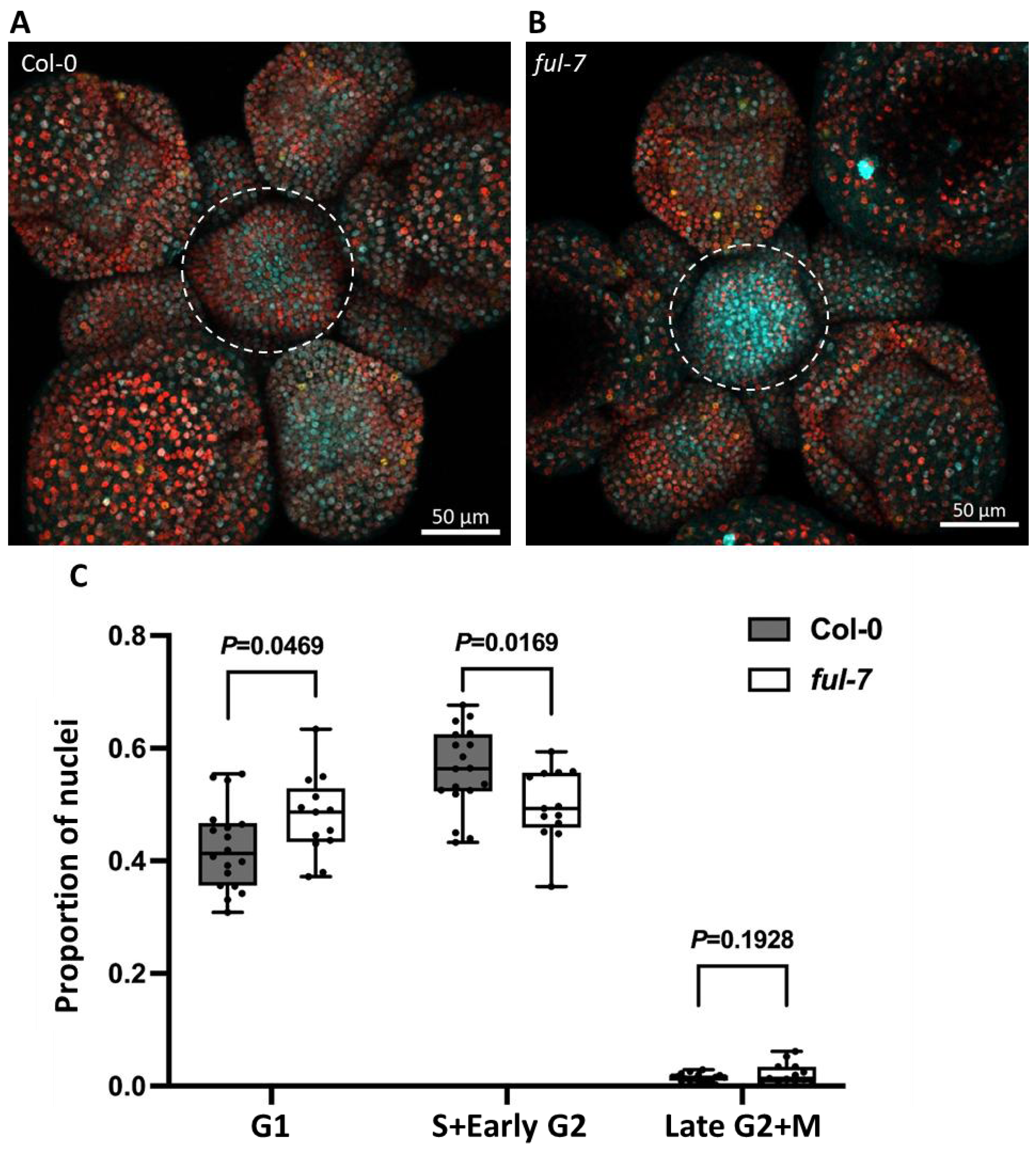
Changes in cell cycle phases of inflorescence meristem (IM) between Col-0 and *ful-7*. **(A-B)**, Confocal image examples of IM with cell cycle markers (PlaCCI) in Col-0 (A) and *ful-7* (B) backgrounds. The white dashed circles indicate the IMs used for calculation in (C). **C)**. The ratio of nuclei detected in each channel to total nuclei. N=18 for Col-0, N=13 for *ful-7*. *P* value was calculated using a two-tailed Student’s *t*-test. G1 phase is represented by the cyan CFP signal, S+Early G2 phase is represented by the red mCherry signal, and late G2+M phase is represented by the yellow YFP signal.

## Discussion

Postembryonal plant development is associated with an intricate coordination of cell cycle, growth and cellular differentiation ^1^. Plant meristems need to replenish and maintain a pool of sufficient undifferentiated cells to allow the continuous development of the plant ^2^. However, little is known about the transcriptomic regulation balancing the maintenance of a pool of undifferentiated cells by cell cycle and their development to differentiate plant organs. The analysis of our snRNA-seq data of different stages of flower development reveals the transcriptomic dynamics happening in the meristematic cells. Our analysis is able to identify two main groups of cells, one pool of cells which transcriptome indicates that belong to different stages of the cell cycle, and another pool of cells which transcriptome indicates that belong to different stages of flower development. This allows us to identify the FUL-KRP2 regulation as an important regulatory module on the decision of the cell of continuing dividing to replenish the pool of undifferentiated cells or to transition to advanced stages of flower development. An interesting hypothesis is that the FUL-KRP2 module is needed to exit the replenishment and maintenance of undifferentiated cells in the IM/FM. We observed that the regulation of KRP2 by FUL is located in the inner regions of the IM and FM (Figure 3, Supplementary Figure S8-9) which may indicate the identity of these cells as undifferentiated stem cells. Previous studies have shown the importance of the cell cycle in the maintenance of pluripotency during flower development ^58^. It has been proposed that the cell-cycle dilution of H3K27me3 in the *KNUCKLES* (*KNU*) promoter, and possible other genes, is needed for a timely floral meristem termination ^59^. This cell-cycle dependent effect was linked to KRPs activities, as the expression of KNU in their system is affected by the *krp1 krp2 krp3 krp4 krp7* quintuple mutant. It is likely that KRP2 plays an important role on this as overexpression of KRP2 leads to the rapid consumption of stem cells in the columella stem region of the root ^60^. Interestingly, FUL has been also implicated in meristem determinacy and termination ^19^, it was proposed that this is an indirect effect achieved by the regulation of AP2 by FUL that in return will regulate the meristem identity marker WUSCHEL (WUS). Taking our results, it is tempting to speculate that the traditional flower development transcription factor FUL is able to influence the start of meristem differentiation and, at the same time, to initiate the count down for meristem termination via the activation of KRP2. In this way, the two main trajectories observed in Figure 2A will correspond to: 1) the replenishment of the pool of undifferentiated cells of the meristem, where the main transcriptomic activity is related with cell cycle regulation and no differentiation programs are active. We denoted this trajectory as “cell cycle trajectory” in Figure 2A. 2) the start of differentiation of the cells, where the transcriptome variation is strongly affected by the differentiation programs, and not so strongly by the cell cycle. We denote this trajectory as “developmental trajectory” in Figure 2A.

Consistent FUL binding to downstream regions in several KRP genes, including *KRP2*, suggests a potentially conserved regulatory mechanism. While downstream binding by MADS-domain transcription factors has been observed previously (Kaufmann et al., *Science* 2010), its role in gene activation remains unclear. One possibility is that such sites facilitate chromatin looping, bringing distal regions into proximity with core promoters. Alternatively, FUL may recruit co-activators or chromatin remodelers that support transcriptional activation from downstream positions. These findings point to a potentially broader regulatory role for MIKC-type MADS-box proteins that warrants further investigation.

We showed that the FUL-KRP2 module is one regulatory point for cell cycle transition in IM/flower meristems. However, the regulation of this process is likely much more complex. Our analysis indicates that FUL also binds genomic regions (Supplementary Figure S6) near other *KRP*s, and known regulators of KRPs (DELLA proteins) ^9^. We predicted that other regulatory factors such as SEP2, ERF74 and KAN2 may also regulate KRP2. In fact, the transgenic KRP2-GFP reporter lines in the *ful-7* mutant background could not completely abolish the KRP2-GFP signal in the meristem (Figure 3). This suggests that there may be other factors regulating *KRP2* expression. Moreover, from our previous study ^17^, FUL binds to two *DELLA* genes (*RGA1* and *RGL2*) and represses the expression of *RGL2* (Supplementary Figure S6) which may also affect KRP2 expression. It has been reported that FUL is also able to control meristem determinacy via the regulation of AP2 which in return regulates the meristem determinacy marker WUS ^19^. Future studies need to reveal the importance of each of these regulations controlling meristem maintenance and differentiation.

KRP2 is reported to control the onset of endoreduplication. In our study we were not able to separate the mitotic cell cycle from the endoreplication cycle. In the shoot apical meristem only euploid cells are observed ^61^, demonstrating that only mitotic division happens on these tissues. They propose that repression of the endoreduplication in the meristem could be a mechanism to ensure genetic stability. They also notice that coinciding with the maturation of the meristematic cell, endoreduplication started to be observed. This indicates that endoreplication is associated with the onset of cell differentiation. Other studies in plants also relate the onset of differentiation with endoreduplication programs ^62,63^. Recently, it has been proposed that endoreduplication is not so rare in mammals, and that is a result from a self-limiting mechanism coordinating cell multiplication with differentiation. In this model, the limit to cell multiplication is imposed by the DNA damage caused by replication stress of many rounds of cell cycle, this DNA damage will trigger the G2/M checkpoint and lead to terminal differentiation which will be associated to endoreduplication.

### Conclusions

Our work illustrates how time-series snRNA-seq experiments can be used to study the regulatory switches between plant developmental programs. In particular, leveraging the system for synchronized floral induction (*pAP1:AP1-GR ap1-1 cal-1)* and in combination with snRNA-seq experiments at different developmental points, we could characterize the transcriptomic changes at the bifurcation point between cell cycle and cell differentiation programs. We computationally identify the activation of KRP2 by FUL as a key point to start switching off the cell cycle program and to continue plant developmental programs during early flower development. We validate this result by quantifying the change in proportion of cell cycle stages in flower meristems of *ful* versus wild-type plants. As expected when FUL controls G2 to M transition, we observed an decrease in the proportion of early G2 cells when in *ful* plants.

## Methods

### Plant Material

All seeds were sterilized and grown on soil, then cold stratified at 4℃in the dark for two days. After cold treatment, all plants were moved into a growth chamber and grown under long-day conditions (16 hours of light, 8 hours of darkness) at around 22 ℃with 60% humidity, and watered every two days.

For plants containing plant cell cycle indicator (PlaCCI) construct, which includes fluorescent sensors, wild-type *Arabidopsis thaliana* Col-0 with PlaCCI was obtained from ^57^. Then, after the removal of stamens from *ful-7* unopened flowers, the stamen of the PlaCCI labeled Col-0 was used to pollinate the stigma of the emasculated *ful-7* flowers. Selected by antibiotic kanamycin and genotyping, the T3 homozygous seeds of *ful-7* with PlaCCI were collected for confocal scanning.

For the KRP2-GFP plants, two constructs with different lengths of downstream regulatory region of *KRP2* were transformed into both Col-0 and *ful-7* plants. The two constructs were initially generated in the pCR8/GW/TOPO entry vector using the In-Fusion HD Cloning Plus (Clontech) kit, both constructs contain the *KRP2* genomic fragment, including 3.6 kb of promoter region and coding sequence of KRP2 without the stop codon, fused with eGFP by a linker sequence. After the GFP sequence, the constructs were ligated either with the downstream 900 bp genomic fragment following the stop codon of *KRP2* or with the extended 1,686 bp genomic fragment, which includes the FUL binding region ^17^. Based on the appearance of the FUL binding region, the constructs were named pKRP2:KRP2-GFP:FUL_peak and pKRP2:KRP2-GFP:w/o_peak. After the LR reactions using Gateway™ LR Clonase™ II Enzyme Mix (Thermo Fisher), the target constructs were transformed into the modified destination vector pMDC32 in which the 35S promoter was removed. Afterwards, we used the electroporation method to transform the terminal vectors into *Agrobacterium*, and transform them into Col-0 and *ful-7* plants by dipping unopened buds into the *Agrobacterium* cell suspension with 5% sucrose and 0.05% Silwet L-77 ^64^. Two transfections a week apart were performed. After selection by genotyping, the T2 plants of positive lines were used for confocal scanning. All primers used in the cloning and genotyping process are listed in Supplementary Table S5.

### Preparation of single-nucleus RNA-seq libraries

*pAP1:AP1-GR ap1-1 cal-1* plants ^65^ were daily induced by Dex solution (2 μM Dexamethasone and 0.016% Silwet L-77) after bolting 0.5 cm to 3 cm tall. Inflorescences from the main stem were collected before DEX-induction (0 day), and after 2, 4, and 8 days of the first induction. Around 20 inflorescences were collected for each sample and used for nuclei isolation.

Nuclei were isolated as previously described ^29^. Briefly, frozen inflorescences were crushed into small pieces manually with pestle and mortar and transferred to a gentleMACS M tube. Samples were then dissociated by gentleMACS in Honda buffer (2.5% Ficoll 400, 5% Dextran T40, 0.4 M sucrose, 10 mM MgCl2, 1 µM DTT, 0.5% Triton X-100, 0.4 U/ µl RiboLock RNase inhibitor and cOmplete^TM^ protease inhibitor). The homogenized plant tissue was filtered through a 70 µm strainer and nuclei were pelleted at 1000 x*g* for 6 min at 4°C. Most Honda buffer was removed and the nuclei pellet was resuspended in the remaining 500 µl Honda buffer. Nuclei were filtered again through a 35 µm strainer and stained with 2∼3 µM DAPI. Single and intact nuclei were sorted using a BD FACS Aria III gating on the DAPI signal.

Single nuclei libraries were prepared using the SMARTer ICELL8 single-cell systems as previously described ^29^. Nuclei were double stained with NucBlue and dispensed into a barcoded SMARTer ICELL8 30 DE Chip by the NanoDispenser. Single nuclei wells were selected using the ICELL8 Imaging Station manually with the support of CellSelect software. 1030, 1320, 837 and 1071 nuclei were selected for library preparation for 0 day, 2 day, 4 day, and 8 day, respectively RT-PCR reaction was done with the SMARTer ICELL8 30 DE Reagent Kit and cDNA was pooled from the chip and concentrated using the Zymo DNA Clean & Concentrator kit. cDNA was then used to construct the DNA libraries using the Illumina Nextera XT kit, followed by AMPure XP bead purification. A Qubit dsDNA HS Assay Kit and a KAPA Library Quantification Kit were used for library DNA quantification, and the Agilent High Sensitivity D1000 ScreenTape Assay was used for library DNA size assessment.

### Single-nucleus RNA-seq data analysis

Raw sequencing files were demultiplexed and the FASTQ files were generated using Illumina bcl2fastq software (v2.20.0). At the moment of the analysis of the data, the associated ICELL8 mapped analysis pipeline (demuxer and analyzer v0.92) didn’t allow to count mapped reads per gene in a strand-specific manner. Therefore, after removing Illumina adapters using Trimmomatic (V0.32), reads were mapped to the Arabidopsis TAIR10 genome using STARseq with parameters: *alignIntronMin*: 20; *alignIntronMax*: 10000; *outFilterMismatchNoverReadLmax*: 0.04; *outFilterMismatchNmax*: 999; *alignSJoverhangMin*: 8; *alignSJDBoverhangMin*: 1; *sjdbScore*: 1. Later, the FeatureCounts software (v2.0.1) was used with parameters: *--primary -R CORE -F GTF -O -s -Q 0 -t gene*. A Python script was used to generate the number of read counts per gene and barcode. Next, only barcodes present in the pre-defined list of barcode sequences included in all Takara Bio NGS kits were considered. Barcodes representing positive or negative control were removed. All samples were combined in one matrix. Genes encoded in the organelles were removed. Only genes with more than 10 read counts in at least 15 barcodes (among all samples) were retained. Nuclei with (i) less than 10, 000 reads, (ii) less than 500 genes containing 10 reads, or (iii) at least one gene covering more than 10% of the reads of a particular nucleus were filtered out. Next, dimensional reduction and clustering were performed using LIGER (rliger v1.0.0), using default parameters except *k=15, lambda=15, min_dist=0.1 and n_neighbors=10*. Clusters identified as meristematic or belonging to sepal primordia were reanalyzed using MONOCLE3 (Cao et al., 2019). We use MONOCLE3 instead of LIGER at this step because the former allowed us to estimate pseudo-time. We used the top 75 marker genes for each meristematic cluster identified with LIGER to run MONOCLE3’s preprocess_cds function with parameter *num_dim = 10, norm_method=“log”*. For the alignment of different samples the function *align_cds* from the package MONOCLE3 was used. The variable number of genes detected and coverage was used in the *residual_model_formula_str* parameter to remove batch effects. For the function *reduce_dimension* we used set parameters to *max_components=2, umap.n_neighbors = 15, cores=1, umap.min_dist=10^-5^, umap.metric = “cosine”, umap.fast_sgd = FALSE*. For the function *cluster_cells* we used the parameters *k=20, cluster_method=“leiden”, resolution=10^-2^, num_iter = 1500*. MONOCLE 3 allowed us to estimate the pseudo-time associated with each cell and their trajectories with the function *learn_graph* using default parameters except: *Euclidean_distance_ratio*=2. We used the spatial autocorrelation analysis called Moran’s I test implemented in the *graph_test* of Monocle3 to test the expression of which cell cycle genes correlate with the two trajectories identified in our snRNA-seq data. For this, we focus our analysis on clusters 6, 2, 7, 15, and 4 which are located around the bifurcation point. We only analyze cell cycle genes as defined in the Supplemental Dataset 1 from Boruc *et al*., 2010 ^50^.

Gene regulatory network was estimated using GENIE3 with default parameters except *nTrees* parameter which was set to 10000 and for the list of regulators (parameter *regulators*) only genes designed as transcription factors in the Plant Transcription Factor Database ^66^ were used. Gene expression was standardized to mean 0 standard deviation 1 before running GENIE3.

### Confocal analysis for KRP2-GFP plants

For KRP2-GFP detection, plants were grown till 1∼3 flowers opened (within one week after bolting) and imaged at 40x on a Zeiss LSM 800 confocal laser scanning microscope using Zeiss 40x water immersion objective (W Plan-Apochromat 40x). Before imaging, the old floral buds were carefully removed, and the center most part was kept and stained for 5 minutes by 1 mg/mL propidium iodide (PI). After being shortly rinsed by water, the sample tissue was used for imaging. One track was used with 488 nm and 561 nm laser to excite the PI and eGFP fluorescences, and emission was collected from 595 nm to 617 nm for PI signal and from 410 nm to 532 nm for eGFP signal. To get better cell segmentation in further analysis, we used the optimal 0.66 µm interval between stacks.

After imaging, the raw czi. files were open in Fiji, and PI and eGFP channels were split and saved as tif. files respectively. The following expression analysis was done by MorphoGraphX (MGX) software ^67^. For cell segmentation, PI tif. files were load in MGX and processed as follows:

1. Stack/Filters/Gaussian Blur Stack: 0.3x0.3x0.3 µm^3^
2. Stack/ITK/Segmentation/ITK Watershed Auto Seeded: Level=600
3. Select the “Delete label in volume” tool and press “Alt” when clicking the generated box.
4. Select the “Delete label in volume” or “Voxel Edit” tool to remove the cells outside IM.
5. Mesh/Creation/Marching Cubes 3D: Cube size=1.0 µm
6. Mesh 1/Save

Next, *Work* and *Labels* in stack 1 were unselected, and load the eGFP tif. file as *Main* figure. Then, to calculate the eGFP signal intensity for the segmented cells, we projected the eGFP signal onto the cell mesh by following steps:

1. Mesh/Signal/Project Signal: Use absolute= Yes, Min Dist=0, Max Dist=3 µm
2. Mesh/Heap Map/Measures/Signal/Signal Total
3. Mesh/Heap Map/Heap Map Save

To capture the signal for each cell, we projected the signals in the 3 µm radius range from the cell mech surface according to the general meristem nuclei size studied before ^68^. Then, the saved heatmap file contains the absolute value of eGFP signal obtained by confocal microscopy for each segmented cell, and it will be used for the statistical analysis.

### Cell cycle proportion measurement of inflorescence meristems

Main inflorescence stems were cut from plants when 1 ∼3 flowers opened and imaged at 40x on a Zeiss LSM 800 confocal laser scanning microscope. To cover each nucleus, scanning intervals were 3 µm between stacks according to the meristem nuclei size studied before ^68^. We used three individual tracks to collect the fluorescence signal of the three fluorescent indicators in the single PlaCCI construct. mCherry was excited with a 561 nm laser, and the emission was collected from 400 nm to 630 nm. YFP was excited with a 488 nm laser, and the emission was collected from 520 to 580 nm. CFP and ChloA (as the IM shape navigator) was excited with a 405 nm laser, and the emission was collected from 410 to 520 nm for CFP and from 650 to 700 for ChloA. In Fiji, IMs were chopped out for analysis and individual fluorescence channels were split and saved as tif. files.

Raw z-stack tif. files of each of the three reporter channels were processed in MGX with the same setting as follows (adapted from Vijayan *et al*., 2021^69^):

1. Stack/Filters/Bright Darken: 3
2. Stack/Filters/Gaussian Blur Stack: 0.2x0.2x0.2 µm^3^ (apply two times)
3. Stack/Segmentation/Local Maxima: x/y/z radius=1.8 µm; threshold=15000
4. Mesh/Creation/Mesh From Local Maxima: radius=1.5 µm
5. Mesh/Heat Map/Analysis/Cell Analysis 3D
6. Mesh/Attributes/Save to CSV Extended

These processes generated comparable nuclei numbers of different cell cycle phases for Col-0 and *ful-7*. As identification of PlaCCI for dividing cells, pCDT1a::CDT1a-eCFP indicates the G1 cells, pHTR13::HTR13-mCherry indicates S+early G2 cells, and pCYCB1;1::NCYCB1;1-YFP represents to late G2+M cells ^57^. Thus, the proportion of different cell phases in IM was calculated as dividing the nuclei number from individual corresponding channels by the summed nuclei number of all channels.

### Data availability

The snRNA-seq data is available GEO Omnibus with id GSE255880.

## Author Contributions and Acknowledgments

J.M.M. and K.K. designed the project. P.C. and X.X. performed experiments. J.M.M., C.S., and K.K. supervised the experiments and contributed to data interpretation. J.M.M. performed data analysis for sequencing results, and P.C. performed the data analysis for microscope images. R.S., B.D and C.G. provided material and discussed results. P.C. and J.M.M. wrote the manuscripts with input from all authors.

This work was supported by grant PID2021-123319NB-I00 (Ministerio de Ciencia e Innovacion, Spain) to C.G.. R.S. is funded by BBSRC grant BB/X01102X/1 and UKRI Frontier Research Grant EP/X034550/1. K.K. wishes to thank the DFG for funding (grant numbers 458750707, 512328399, 438774542, 546593285).

## Table legends

**Supplementary Table S1. Marker genes of the 11 clusters shown in Figure 1**.

**Supplementary Table S2. Marker genes of the 18 clusters shown in Figure 2**.

**Supplementary Table S3. Estimated gene regulators of KRP2 in cluster 15**. Gene expression regulators of KRP2 estimated by GENIE3.

**Supplementary Table S4. FUL’s ChIP binding scores and regulations on *KRPs* and *DELLAs* in IM tissues**. Scores of FUL binding peaks within ±3 kb of *KRP* and *DELLA* genes, detected by ChIP-seq of IM tissues. Gene expression changes of *KRP* and *DELLA* genes (*ful-7* vs Col-0) in IM tissues. Data was obtained from ^17^

**Supplementary Table S5. List of primers used.**

