## supplementary figures for "The role of FRUITFULL controlling cell cycle during early flower development revealed by time series snRNA-seq experiments"

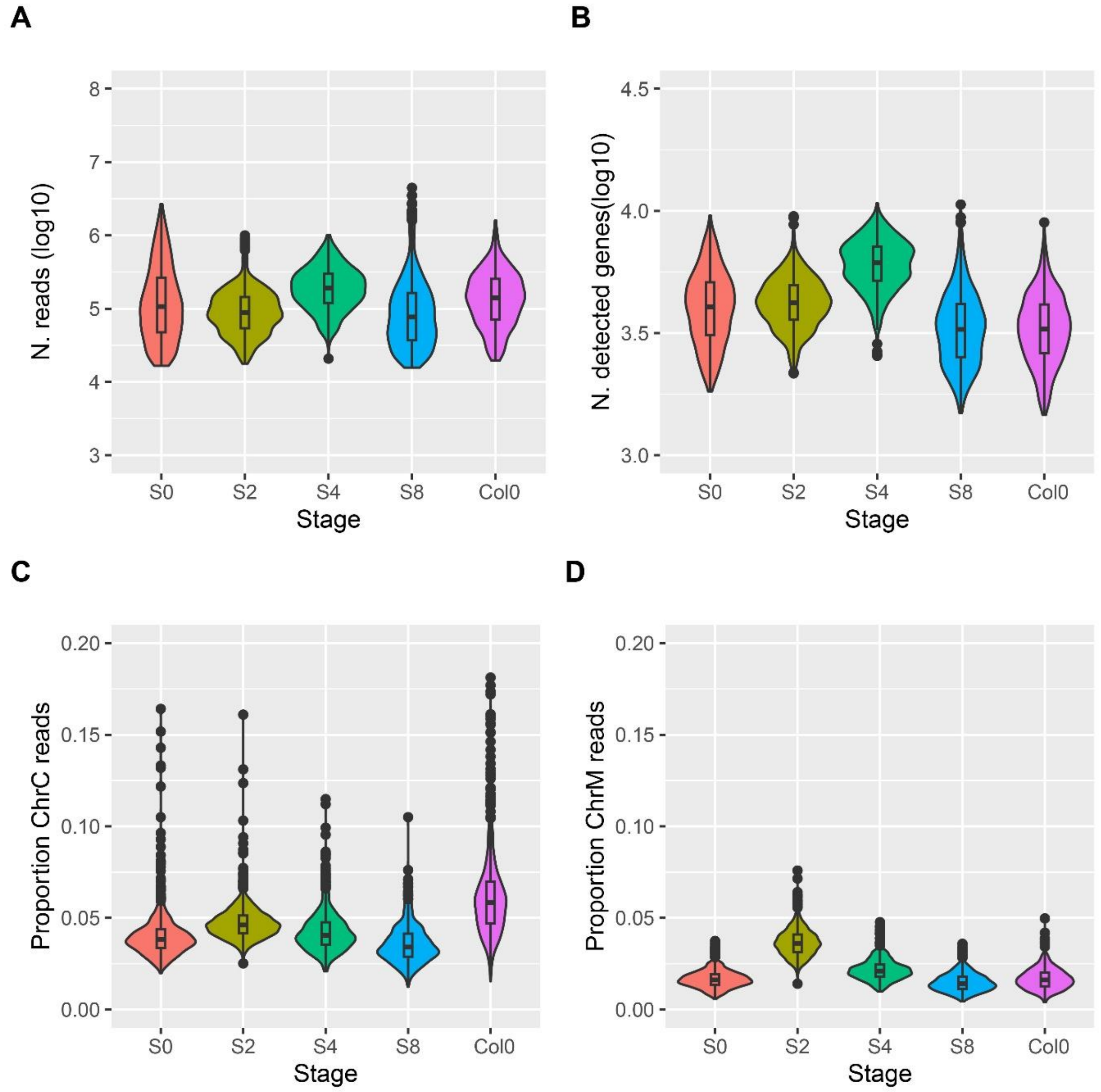


**Supplementary Figure S1. Quality control of the snRNA-seq experiment. A)** Distribution of the number of reads mapped per nucleus. **B)** Distribution of the number of detected genes per nucleus. **C)** Proportion of mapped reads located in the chloroplast genome per nucleus. **D)** Proportion of mapped reads located in the mitochondrial genome per nucleus.

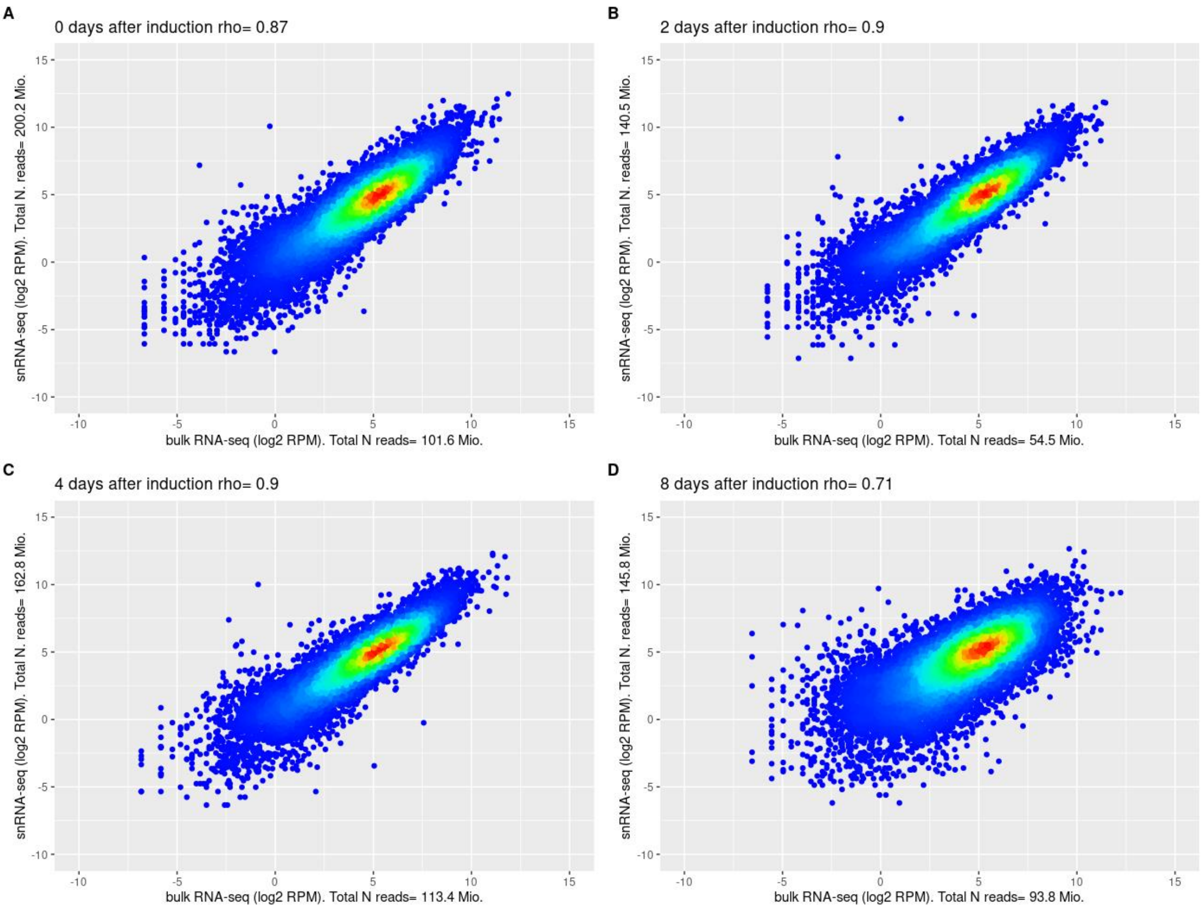


**Supplementary Figure S2. Reproducibility between snRNA-seq and bulk RNA-seq**. The scatterplot shows the high reproducibility of the different snRNA-seq datasets against bulk RNA-seq data collected at the same stage and genotype (Yan et al., 2019) (A-D). For snRNA-seq, read counts per protein-coding gene were summed across all transcriptomes. For bulk RNA-seq, read counts per protein-coding gene were summed across all biological replicates. Read counts per gene were normalized by the total number of mapped reads and multiplied by $10^6$.

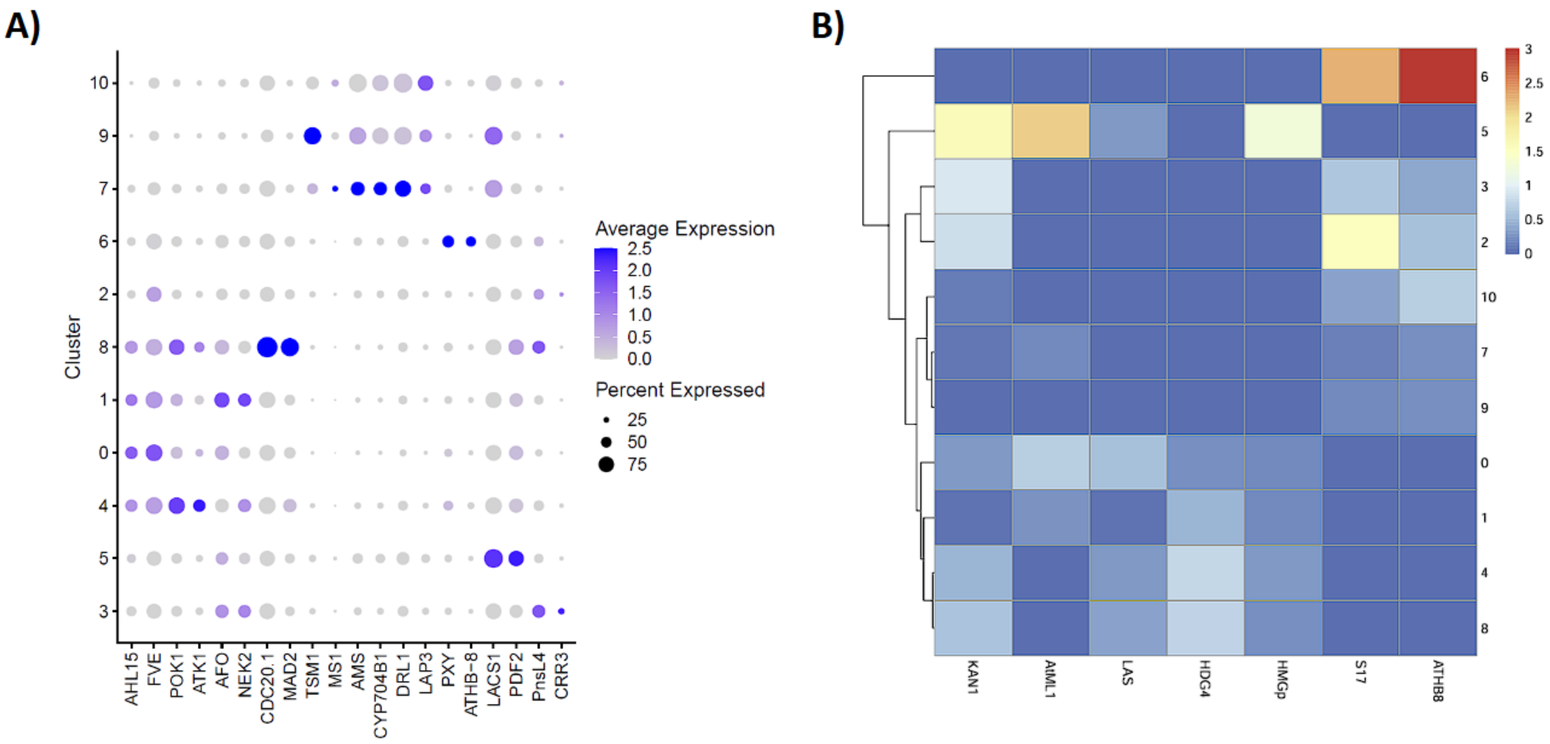


**Supplemental Figure S3. Expression and coexpression of snRNA-seq marker genes. A)** Dotplot showing the average expression and percent of cells expressing particular genes mentioned in the main text to help cluster annotation. **B)** Heatmap showing the Pearson correlation between the average expression of each cluster (rows) and selected bulk RNA-seq data from (Yadav et al., 2014) (columns). To calculate the correlation only genes identified as top50 marker genes for each cluster were used.

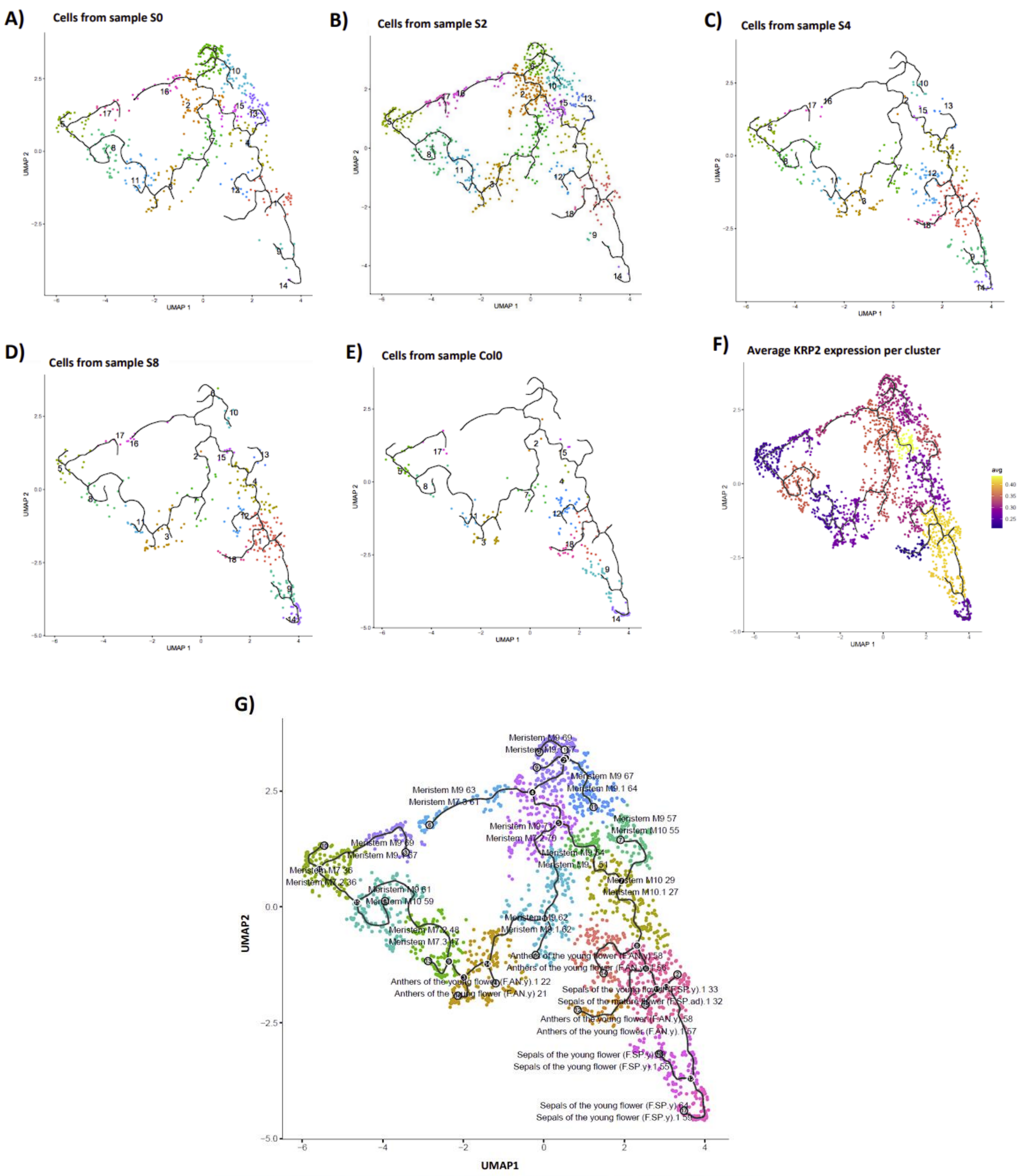


**Supplementary Figure S4. Distribution of snRNA-seq samples across the UMAP plot**. The UMAP plot shown in Figure 2a is plotted for each snRNA-seq dataset separately. Points are color-coded as Fig 2a to indicate cluster groups. **A**) it shows only transcriptomes from the DEX-uninduced sample (S0), **B**) from the S2 sample, **C**) from the S4, **D**) from the S8 sample, and **E**) from the Col0 sample. **F**) Average expression of KRP2 among the transcriptomes of each UMAP cluster. **G)** Annotation of each cluster depending on its expression correlation with bulk RNA-seq (TraVa, flower tissues). The correlation was calculated between the average gene expression for each cluster and TraVA developmental stage-specific bulk RNA-seq data. The two TraVa datasets with the highest correlation are reported for each cluster. The last 2 digits represent the correlation coefficient *100.

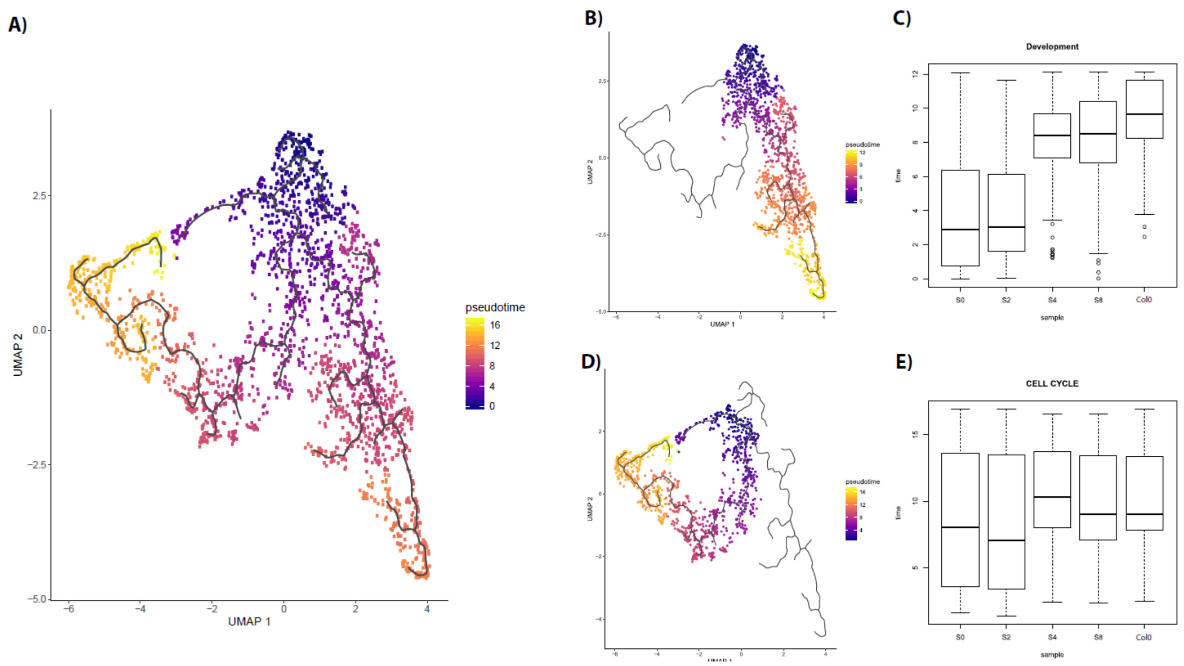


**Supplementary Figure S5. Visualization of pseudo-time estimation in the UMAP plot**. Pseudo-time was estimated for all cells using Monocle3. Cluster 6 was set at pseudo-time 0 as it was the cluster with the highest enrichment in transcriptomes from the snRNA-seq sample S0. **A**) UMAP plot showing the pseudo-time values estimated for each transcriptome. **B**) UMAP plot showing the pseudo-time estimated for each transcriptome of the clusters linked to the development-related trajectory. **C**) Boxplot of pseudo-time values estimated for each transcriptome represented in B) depending from which snRNA-seq sample they belong. The boxplot shows a significant difference in pseudo-time distribution between samples. **D**) UMAP plot showing the pseudo-time values for each transcriptome associated with the cell cycle trajectory. **E**) Boxplot of pseudo-time values estimated for each transcriptome represented in D) depending from which snRNA-seq sample they belong. The boxplot shows no significant differences in pseudo-time distribution between samples.

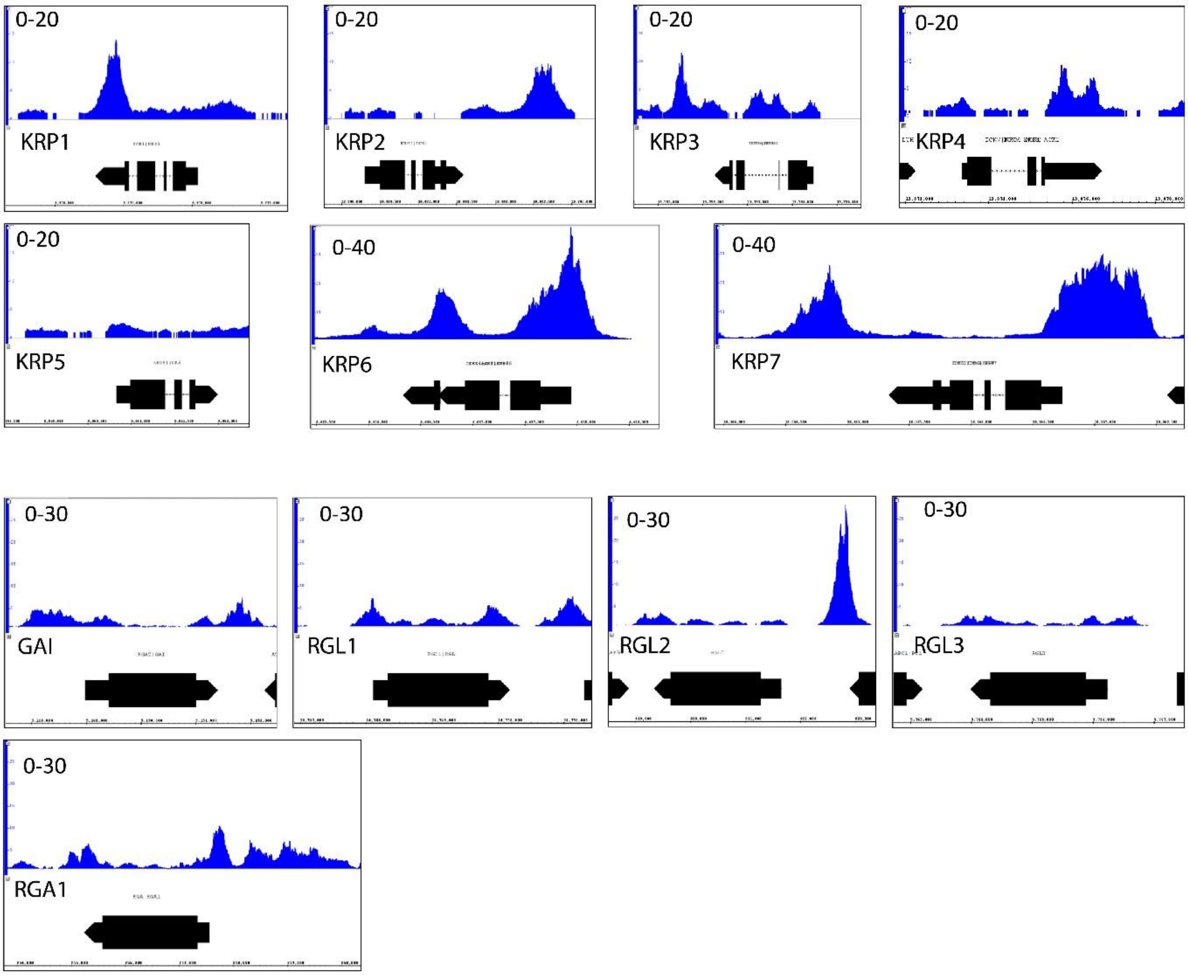


**Supplementary Figure S6. FUL binding patterns in KRP and DELLA protein-coding genes**. Screenshots of the genomic regions around KRP and DELLA protein-coding genes. The screenshots show FUL DNA binding from ChIP-seq experiments (Van Mourik et al., 2023). *KRP* genes, except for *KRP5*, have a significant FUL binding peak (FDR<0.05). For *DELLA* genes, only *RGL2* and *RGA1* have a significant FUL binding peak (FDR<0.05). See Supplementary Table S4 for FDR values.

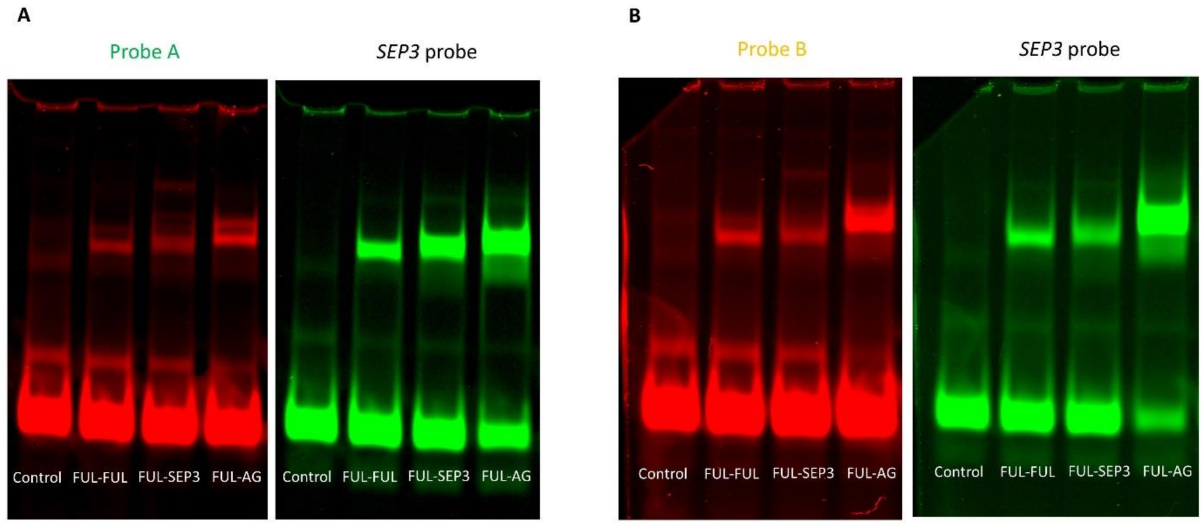


**Supplementary Figure S7. EMSA showing FUL binding to the *KRP2* downstream genomic region**. EMSA results of *KRP2* probes with different FUL dimers. Probe A and B represented the CArG-box A (A) and B (B) sequences located in the *KRP2* downstream region within the FUL binding peak (see Figure 3a). As positive control, a probe called "*SEP3* probe" representing a DNA fragment from *SEPALLATA3* promoter region (a shorter version of "SEP3 wt" from (Smaczniak et al., 2012) was used. The dimer tested were FUL homodimer (FUL) and FUL heterodimers with AGAMOUS (FUL-AG) and SEPALLATA3 (FUL-SEP3).

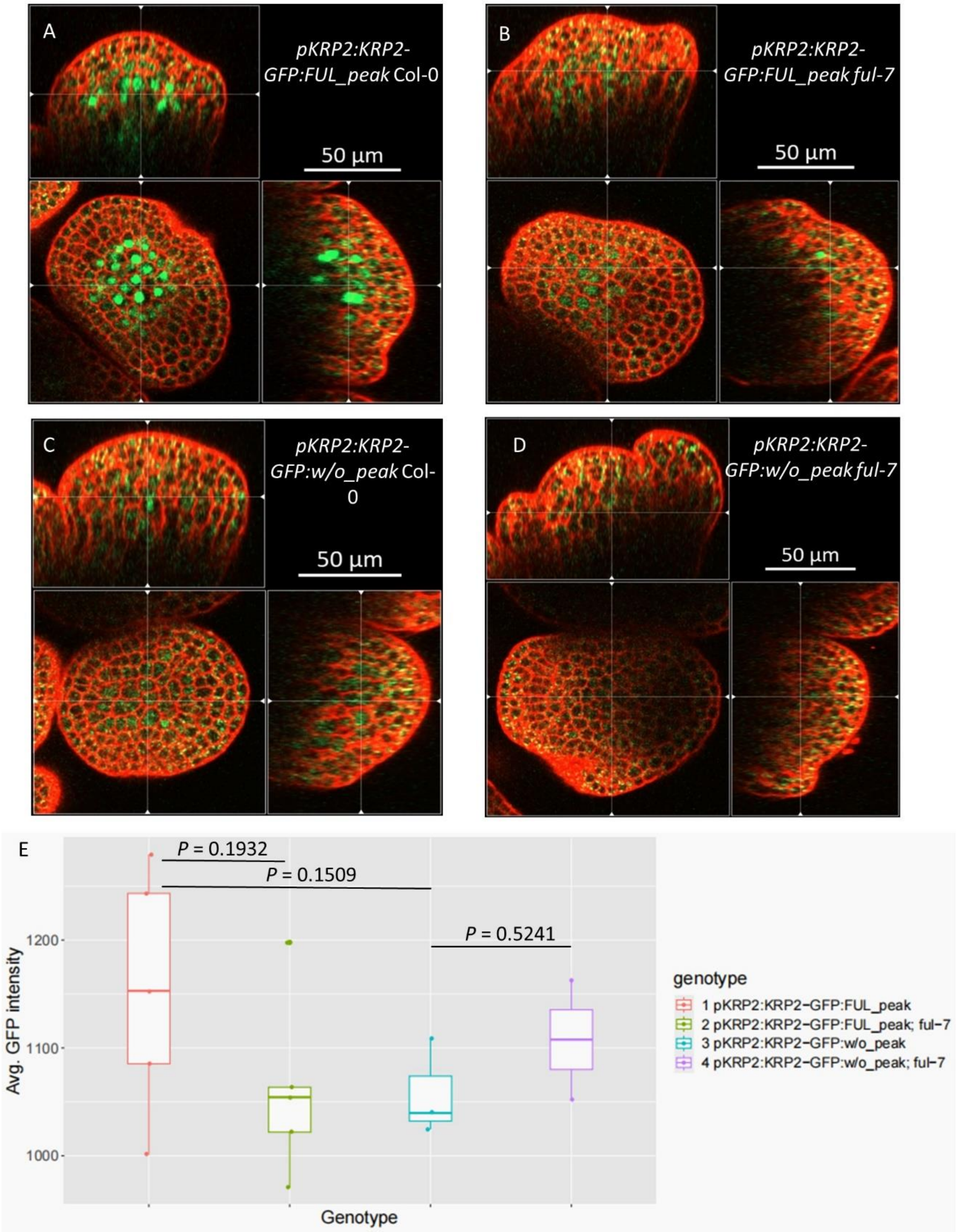


**Supplementary Figure S8. Confocal analysis of KRP2-GFP intensity in stage 3 flowers. A-D)** Confocal images of stage 3 flowers with ortho views. The genotypes are: **A)** *pKRP2:KRP2-GFP:FUL_peak* (N = 5), **B)** *pKRP2:KRP2-GFP:FUL_peak ful-7* (N = 5), **C)** *pKRP2:KRP2-GFP:w/o_peak* (N = 3), **D)** *pKRP2:KRP2-GFP:w/o_peak ful-7* (N = 2). The plants used for scanning are at a similar age with only 1-3 open flowers. **E)** Average GFP intensity of stage 3 flower meristem cells. Bilateral *t*-test was used to test for differences in intensity

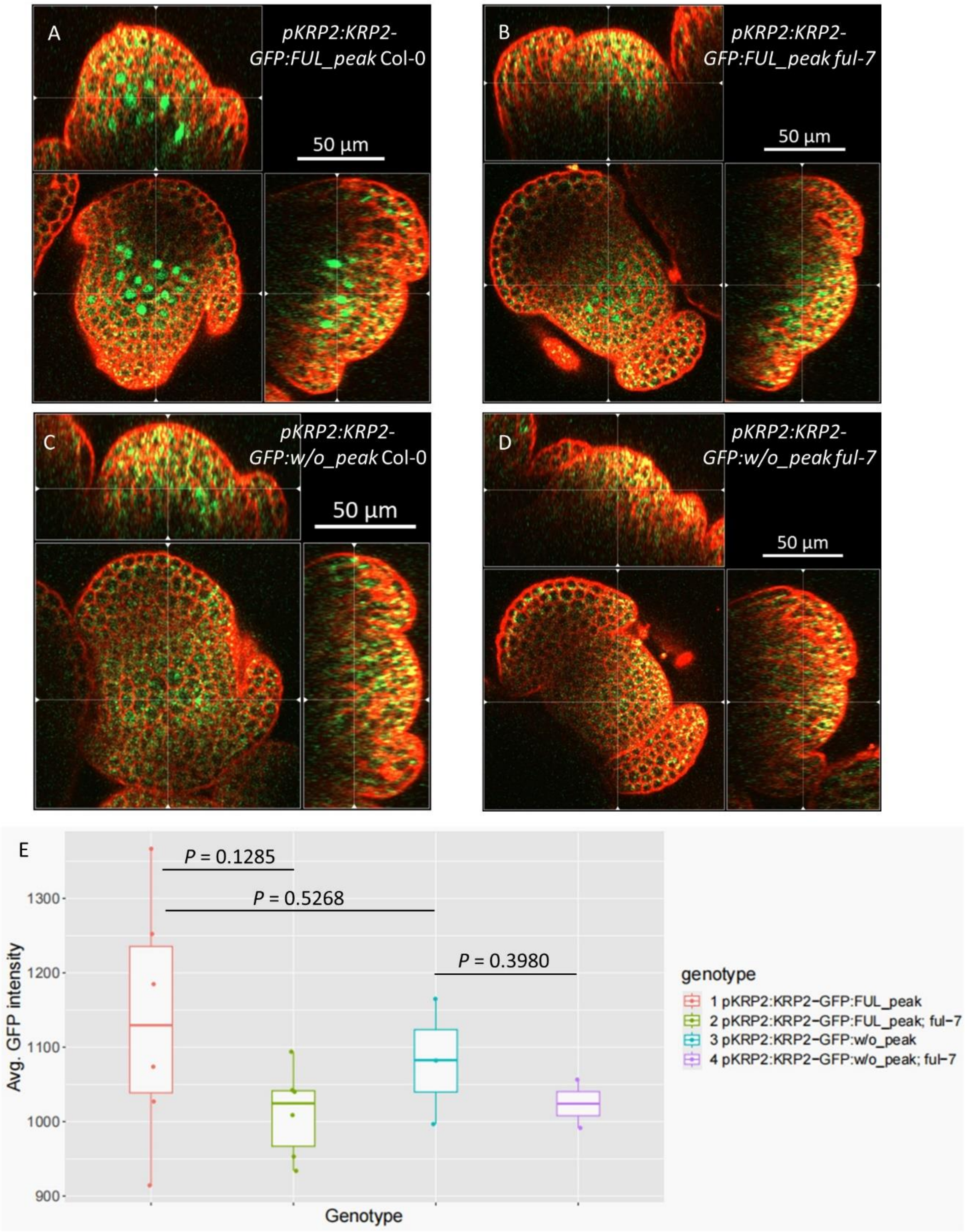


**Supplementary Figure S9. Confocal analysis of KRP2-GFP intensity in stage 4 flowers. A-D)** Confocal images of stage 4 flowers with ortho views. The genotypes are: **A)** *pKRP2:KRP2-GFP:FUL_peak* (N = 6), **B)** *pKRP2:KRP2-GFP:FUL_peak ful-7* (N = 6), **C)** *pKRP2:KRP2-GFP:w/o_peak* (N = 3), **D)** *pKRP2:KRP2-GFP:w/o_peak ful-7* (N = 2). The plants used for scanning are at a similar age with only 1-3 open flowers. **E)** Average GFP intensity of stage 4 flower cells. Bilateral *t*-test showed no significant differences.

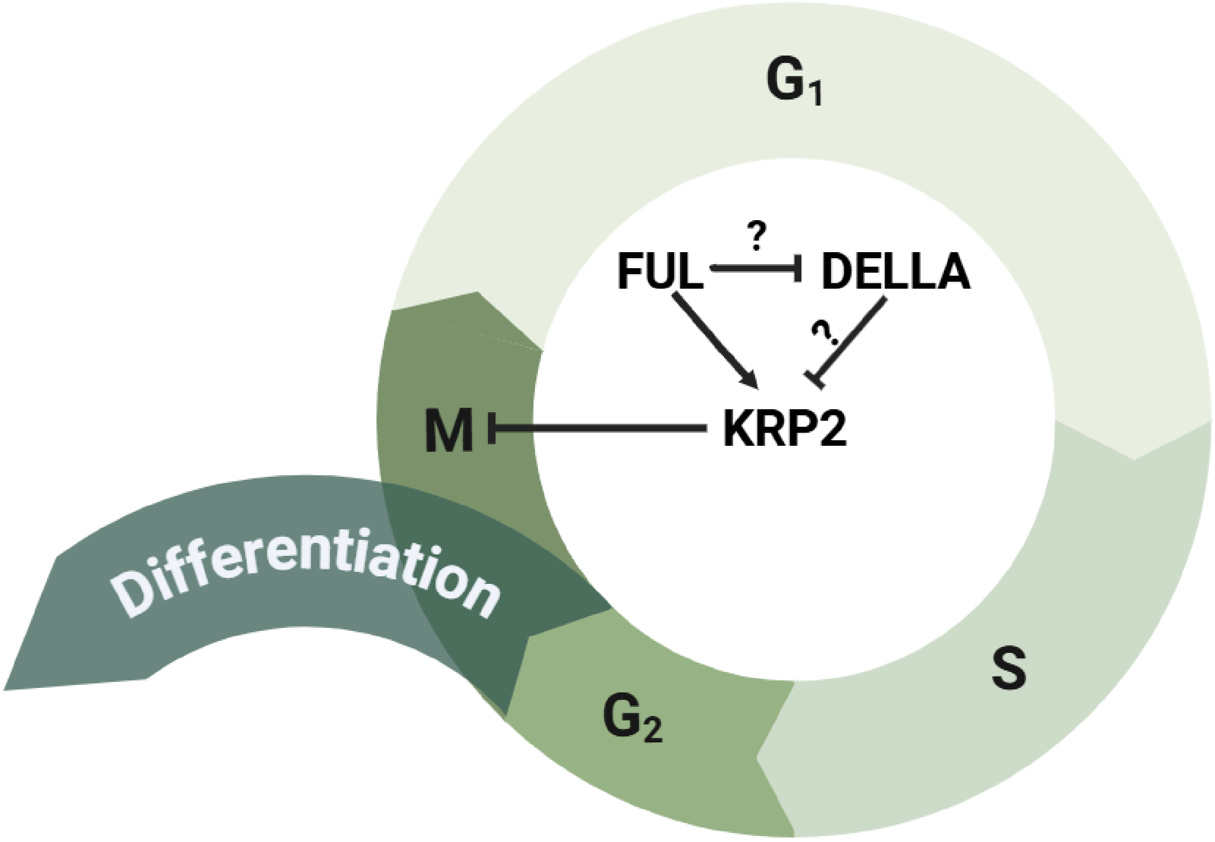


**Supplementary Figure S10. Summary of the FUL-KRP2 regulation.** FUL is able to bind a regulatory region downstream of *KRP2* and regulate *KRP2* expression as validated by *in vivo* and *in vitro* experiments in this study. KRP2 is reported to control G2-to-M transition (see Background section). Our FUL ChIP-seq data also shows that FUL could potentially regulate several DELLA genes (*RGA1* and *RGL2*). These DELLA proteins are involved in the regulation of *KRPs* expression.
